# Adaptive Cluster-Count Autoencoders with Dirichlet Process Priors for Geometry-Aware Single-Cell Representation Learning

**DOI:** 10.64898/2026.03.26.714611

**Authors:** Zeyu Fu

**Affiliations:** State Key Laboratory of Trauma and Chemical Poisoning, Institute of Combined Injury, Chongqing Engineering Research Center for Nanomedicine, College of Preventive Medicine, Army Medical University, Chongqing, China

**Keywords:** single-cell RNA-seq, representation learning, Dirichlet process mixture model, flow matching, autoencoder, latent space, dimensionality reduction

## Abstract

Standard autoencoders for single-cell transcriptomics learn latent spaces whose cluster structure emerges only post hoc through *K*-means or community detection, leaving cluster count and boundary quality uncontrolled during training. Here we ask whether imposing an adaptive nonparametric prior can shift this balance. We equip a feedforward autoen-coder with an online Dirichlet Process Mixture Model (DPMM) prior that refits cluster assignments throughout training and directly regularizes latent compactness and separation. Across 56 scRNA-seq datasets the DPMM prior produces a pronounced *geometry– concordance trade-off* : cluster compactness (ASW) improves by 127% and Davies–Bouldin overlap drops by 47%, but label-recovery metrics decline (NMI −17%, ARI −21%) and downstream *k*NN accuracy falls from 0.784 to 0.725. Wilcoxon signed-rank tests confirm that the geometry gains are significant with large Cliff’s *δ* effects while concordance losses remain bounded and non-significant. A second-stage conditional-flow refinement (DPMM-FM) further improves projection fidelity (DRE 0.751, LSE 0.695, DREX 0.873) at additional concordance cost, revealing a three-tier operating regime: prior-free for label recovery, DPMM for manifold geometry, and DPMM-FM for visualization fidelity. Against 18 external baselines DPMM-Base wins 70.5% of core-metric comparisons (*p*<0.05). Gene Ontology enrichment confirms that geometry-improved latent components recover coherent biological programs. Rather than claiming universal superiority, this study characterizes the operating envelope of nonparametric mixture priors and identifies the task contexts— trajectory analysis, manifold visualization, and program-level annotation—where adaptive geometric structure outweighs label-counting accuracy.

## 1. Introduction

How much geometric structure should a latent space impose during training, and what does that structure cost? Most autoencoder pipelines for single-cell RNA sequencing (scRNA-seq) sidestep this question: the encoder is trained with a reconstruction objective alone, and cluster structure is recovered post hoc by running *K*-means or Leiden [1] on the embedding [2,3]. The resulting latent spaces often achieve high label concordance (NMI, ARI) but poor cluster compactness and boundary quality, because no training signal penalizes geometric diffuseness [4,5]. Recent geometry-aware approaches regularize the latent manifold directly [6], yet none integrate an adaptive, data-driven cluster prior that jointly controls cluster count and boundary quality during representation learning.

An alternative is to impose a structured nonparametric prior that explicitly regularizes the latent manifold. Dirichlet Process Mixture Models (DPMMs) [7,8] are attractive for this role because they do not require the cluster count to be specified in advance: an online Bayesian Gaussian Mixture [9] adaptively partitions the latent space, creating and merging components as the data warrant. A concurrent approach, scDAC [10], couples an autoencoder with a DPMM for adaptive clustering but focuses on concordance optimization rather than characterizing the full geometry–concordance trade-off. More broadly, directly constraining the latent distribution toward a mixture of Gaussians may pull representations away from the configurations that maximize label recovery—a trade-off that has not been systematically characterized on a large single-cell benchmark.

This paper provides that characterization. We equip a standard feedforward autoencoder with a DPMM prior (DPMM-Base) and a further conditional optimal-transport flow refinement stage (DPMM-FM) [11,12], then evaluate a three-model progression—prior-free Pure-AE, DPMM-Base, and DPMM-FM—across 56 scRNA-seq datasets and 41 metrics. Our goal is not to claim that the nonparametric prior is universally superior but to map its operating envelope: *where* geometry gains appear, *what* concordance and downstream-classification costs they carry, and *when* the trade-off is favorable. We find that DPMM-Base produces a 127% ASW improvement and a 47% DAV reduction at the expense of −17% NMI and −21% ARI, defining a regime suited to trajectory analysis [13] and manifold visualization rather than label counting. DPMM-FM extends this profile toward projection fidelity (DRE, LSE, DREX) at additional concordance cost. Together, the three models span a controlled geometry–concordance axis that lets practitioners select the right operating point for their biological question.

## 2. Methods

### 2.1. Model Progression

The experimental design is a controlled ablation along a single axis: latent prior complexity. All three models share the same encoder–decoder backbone; only the prior and optional flow refinement differ.

**Pure-AE** (prior-free baseline) is a standard feedforward autoencoder (encoder layers [256, 128], latent dimension 10) with mean-squared-error reconstruction loss [14]. It serves as the reference for quantifying the contributions of the DPMM prior and flow-matching regularization.

**DPMM-Base** augments Pure-AE with an online Bayesian Gaussian Mixture [9,15] that refits every 10 epochs on latent vectors after a 90% warmup phase. Six DPMM loss options are supported (NLL, KL, energy, Student-*t*, MMD, soft-NLL), enabling flexible cluster-level regularization.

**DPMM-FM** (Flow Matching), our proposed model, augments DPMM-Base with a latent-space conditional optimal-transport flow matching module [11,12] (hidden layers [128, 128], flow weight *λ*_FM_=0.1, noise scale 0.5). The flow head learns a vector field that transports latent samples toward their DPMM-assigned cluster centers via the optimal-transport conditional probability path. This is applied *after* DPMM refitting, smoothing the latent manifold geometry while preserving the Bayesian cluster structure.

All variants share latent dimension 10 and the same encoder–decoder backbone; the key design axis is therefore latent *prior structure* (none vs. adaptive DPMM vs. DPMM + flow refinement), not encoder capacity changes. Figure 1 illustrates the full DPMM-FM architecture.

**Figure 1.**
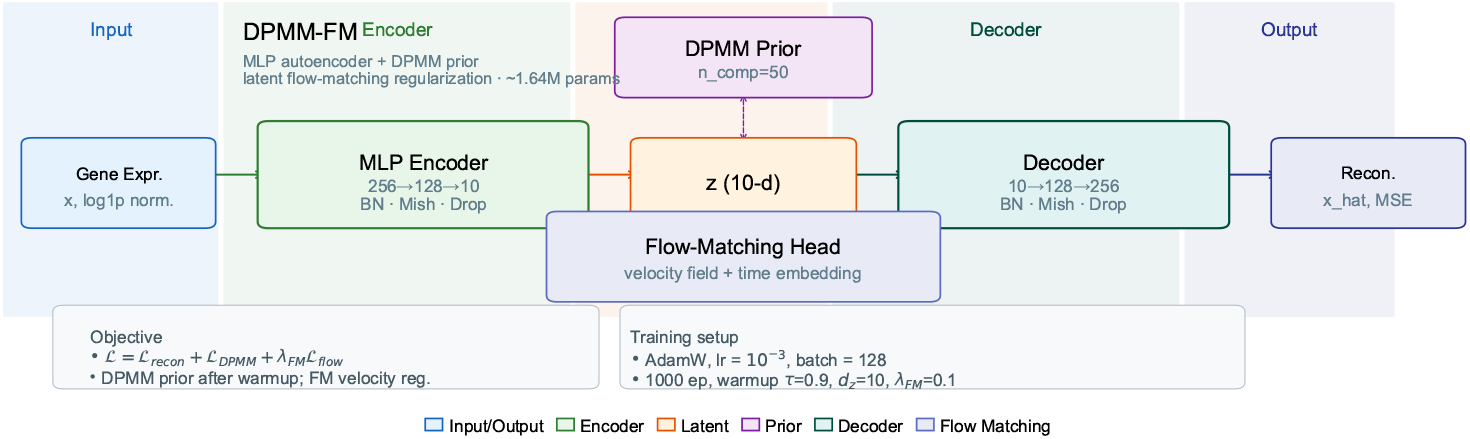
DPMM-FM architecture. A feedforward autoencoder backbone (encoder layers [256, 128], latent dimension 10) with online Bayesian Gaussian Mixture refitting [9] every 10 epochs post-warmup (90%). The optional flow-matching head [11] (hidden [128, 128], *λ*_FM_=0.1) transports latent samples toward DPMM-assigned cluster centers via conditional optimal-transport paths, smoothing the manifold geometry while preserving Bayesian cluster structure.

### 2.2. Training Protocol

Training runs for 1000 epochs with learning rate 10^−3^, batch size 128, and dropout 0.15. The critical hyperparameter is the warmup ratio (0.9): DPMM regularization is withheld during the first 900 epochs so that the autoencoder converges to a stable reconstruction before adaptive mixture forces reshape the latent space. Activating the prior too early risks premature over-partitioning, where the mixture absorbs reconstruction error as spurious components.

### 2.3. Data and Preprocessing

Input features are the top 3000 highly variable genes (HVGs) after library-size normalization and log(1+*x*) transformation [16], subsampled to 3000 cells per dataset for computational uniformity. The benchmark catalogue comprises 56 datasets (16 core cohorts plus 40 additional preprocessed collections) spanning haematopoiesis, neural development, endoderm differentiation, immune activation, and metastatic tissues across human and mouse.

### 2.4. Evaluation Protocol

To characterize the geometry–concordance trade-off, we employ a 41-metric protocol organized into six families. Metrics are grouped by what they measure, and we interpret them asymmetrically: geometry metrics (ASW, DAV) are the primary endpoints for the DPMM prior; concordance metrics (NMI, ARI) serve as cost monitors; projection-fidelity metrics (DRE, LSE, DREX) are downstream diagnostics.

#### Label concordance

NMI (Normalized Mutual Information) [17] and ARI (Adjusted Rand Index) [18], computed by *K*-means clustering (*K* = number of ground-truth labels) on latent vectors.

#### Geometric structure

ASW (Average Silhouette Width) [19], DAV (Davies–Bouldin Index, ↓) [20], CAL (Calinski–Harabasz, ↑) [21], COR (latent correlation, ↑).

#### DRE (Dimensionality Reduction Evaluation)

Applied to both UMAP [22] and t-SNE [23] projections: distance correlation, local and global neighborhood preservation (*Q*_local_, *Q*_global_), and an overall quality composite. 8 sub-metrics total.

#### LSE (Latent Structure Evaluation)

Intrinsic latent space properties: manifold dimensionality, spectral decay rate, participation ratio, anisotropy score, trajectory directionality, noise resilience, and overall quality composite. 7 sub-metrics.

#### DREX (Extended DR Fidelity)

Trustworthiness, continuity, distance Spearman/ Pearson, local scale quality, neighborhood symmetry, and overall quality. 7 sub-metrics.

#### LSEX (Extended Latent Structure)

Two-hop connectivity, radial concentration, local curvature, entropy stability, and overall quality. 5 sub-metrics.

The composite Score used for model ranking is Score = (NMI + ARI + ASW)/3. This metric weights concordance (NMI, ARI) at two-thirds and geometry (ASW) at one-third; projection fidelity metrics (DRE, LSE, DREX) are not captured by the composite.

Because DPMM-AE is designed around adaptive latent partitioning, we interpret ASW and DAV as primary geometry endpoints and NMI/ARI as assignment-concordance checks. DRE/LSE-family metrics are treated as downstream manifold-fidelity diagnostics, especially for assessing the incremental value of the flow-refinement stage.

## 3. Results

### 3.1. Full-Catalogue Refresh

Table 1 shows the mean performance of the DPMM progression across all 56 datasets.

**Table 1.** 56-dataset refresh summary. Mean metric values across 56 datasets. Higher is better except DAV. Composite = (NMI + ARI + ASW) / 3.

| Model | NMI | ARI | ASW | DAV↓ | DRE-UMAP | LSE | Composite |
| --- | --- | --- | --- | --- | --- | --- | --- |
| Pure-AE | **0.609** | **0.406** | 0.165 | 1.624 | 0.568 | 0.451 | 0.393 |
| DPMM-Base | 0.506 | 0.320 | **0.374** | **0.868** | **0.682** | **0.525** | **0.400** |

The catalogue-level pattern is unambiguous: the DPMM prior buys geometry at the expense of label recovery. Pure-AE leads on concordance (NMI 0.609, ARI 0.406) while DPMM-Base leads on geometry (ASW 0.374, DAV 0.868). The improvements are large— ASW rises by 127%, DAV falls by 47%—but so are the costs (−17% NMI, −21% ARI). The composite score (0.400 vs. 0.393) favors DPMM-Base by only 1.8%, a margin driven entirely by the ASW component outweighing concordance losses. This narrow composite margin should not be mistaken for a decisive win; rather, it signals a genuine regime difference. The two models optimize different aspects of latent quality, and the preferred choice depends on whether the downstream task rewards tight clusters (trajectory inference, visualization) or accurate *K*-means label assignment.

### 3.2. Internal Ablation

Figure 2 presents the ablation analysis comparing Pure-AE, DPMM-Base, and DPMM-FM on the 12 core datasets where all three variants were evaluated.

**Figure 2.**
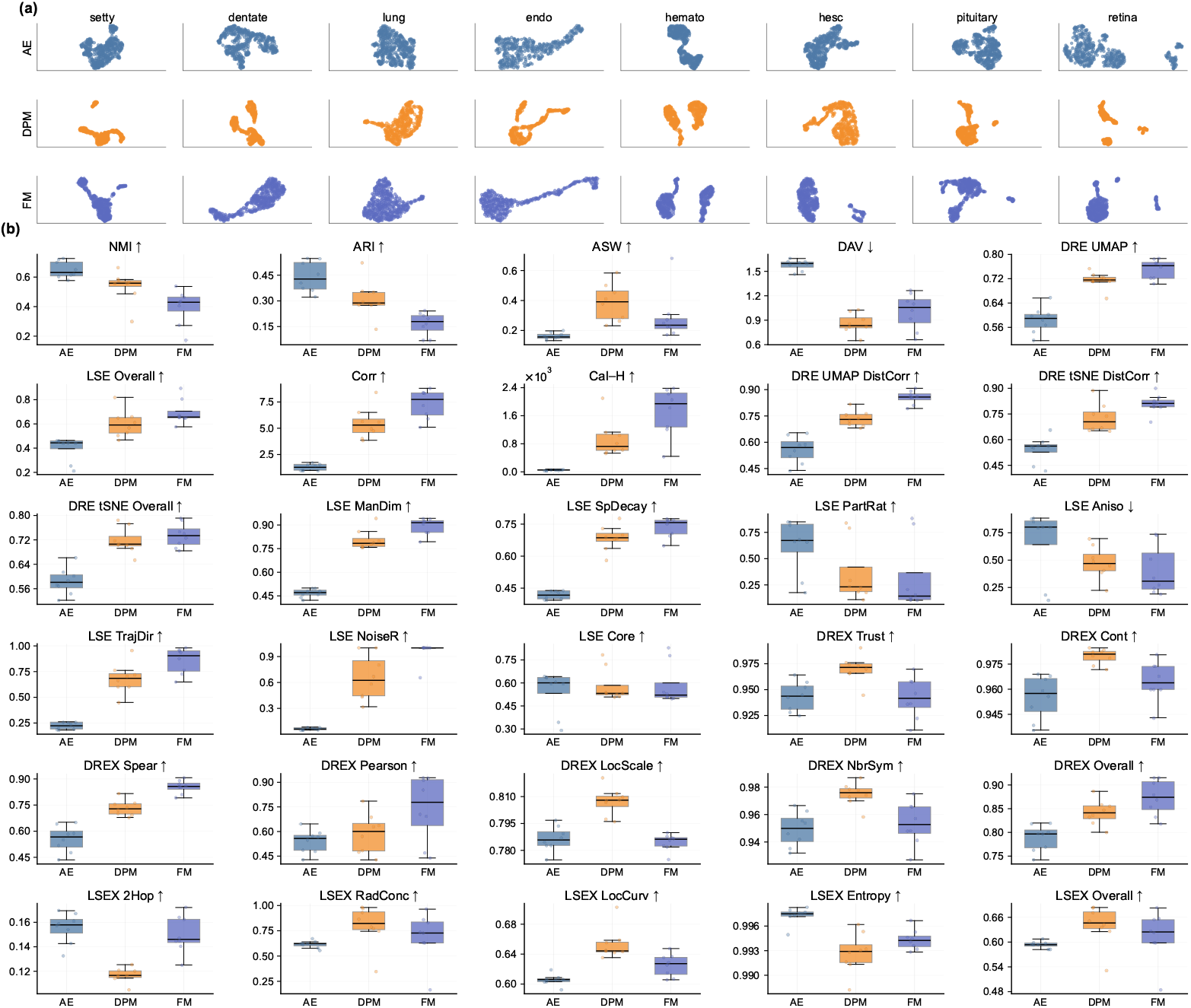
Internal ablation: Pure-AE vs. DPMM-Base vs. DPMM-FM. Panel (a): experimental workflow showing 12 scRNA-seq datasets processed through a controlled three-model ablation (*±*DPMM, *±*flow matching) with 41-metric evaluation. Panel (b): metric boxplot grid across six evaluation families—clustering concordance, DRE-UMAP [22], DRE-tSNE [23], LSE intrinsic, DREX, and LSEX—comparing all model variants.

Table 2 quantifies the three-model comparison on the 12 core datasets.

**Table 2.** Three-model ablation on 12 core datasets. Mean metric values. Best per column in bold. The composite score (NMI+ARI+ASW)/3 weights concordance 2:1 over geometry and does not reflect projection fidelity, where DPMM-FM leads.

| Model | NMI | ARI | ASW | DAV↓ | DRE | LSE | DREX | Score |
| --- | --- | --- | --- | --- | --- | --- | --- | --- |
| Pure-AE | **0.647** | **0.479** | 0.172 | 1.580 | 0.603 | 0.389 | 0.795 | **0.433** |
| DPMM-Base | 0.499 | 0.277 | **0.449** | **0.848** | 0.702 | 0.602 | 0.832 | 0.408 |
| DPMM-FM | 0.397 | 0.165 | 0.288 | 1.005 | **0.751** | **0.695** | **0.873** | 0.283 |

The ablation reveals three distinct operating regimes arranged along a geometry– concordance axis. Pure-AE occupies the concordance end (NMI 0.647, ARI 0.479) with the weakest geometry (ASW 0.172, DAV 1.580). DPMM-Base sits at the geometry end (ASW 0.449, DAV 0.848) with reduced concordance (NMI 0.499, ARI 0.277). DPMM-FM extends the axis toward projection fidelity (DRE 0.751, LSE 0.695, DREX 0.873) at further concordance cost (NMI 0.397, ARI 0.165). Critically, DPMM-FM does *not* dominate DPMM-Base on geometry: its ASW (0.288) and DAV (1.005) are weaker, because the flow field smooths local manifold structure at the expense of tight cluster boundaries. This means the three models are not a simple progression but a Pareto front: Pure-AE for labeling, DPMM-Base for compact clusters, DPMM-FM for smooth low-dimensional projections. The practical recommendation is to select the variant that matches the downstream objective.

### 3.3. Statistical Validation

Table 3 presents pairwise Wilcoxon signed-rank tests [24] confirming which improvements are statistically significant, with effect sizes quantified by Cliff’s *δ* [25].

**Table 3.** Wilcoxon signed-rank tests on core metrics. DPMM-Base vs. Pure-AE across 12 core datasets. W/L/T = wins/losses/ties; Sig. at *α*=0.05.

| Comparison | Metric | W/L/T | $p$-value | Sig. | Cliff’s $\delta$ | Effect |
| --- | --- | --- | --- | --- | --- | --- |
| DPMM-Base vs Pure-AE | ARI | 4/8/0 | 0.924 | No | −0.431 | medium |
| DPMM-Base vs Pure-AE | ASW | 12/0/0 | 2.44e-04 | Yes | 1.000 | large |
| DPMM-Base vs Pure-AE | DAV | 12/0/0 | 2.44e-04 | Yes | 1.000 | large |
| DPMM-Base vs Pure-AE | NMI | 2/10/0 | 0.999 | No | −0.639 | large |

ASW and DAV improvements are significant (*p* = 2.44 *×* 10^−4^, Cliff’s *δ* = 1.00, large effect) across the 12 core datasets: the DPMM prior wins on geometry in every dataset. NMI and ARI losses do not reach statistical significance (*p* = 0.999 and *p* = 0.924, respectively) despite substantial effect sizes (Cliff’s *δ* = −0.639 for NMI, −0.431 for ARI), indicating that while concordance drops are consistent in direction, they are not uniformly large enough to overcome the signed-rank threshold. The asymmetry is informative: geometry gains are universal (12/0 wins), while concordance losses are predominant but not absolute (NMI: 2 wins / 10 losses; ARI: 4 wins / 8 losses). Some datasets benefit on both axes; the majority do not.

### 3.4. Sensitivity Analysis and Training Dynamics

Figure 3 shows hyperparameter sensitivity for DPMM-Base and DPMM-FM alongside training dynamics for the three model variants.

**Figure 3.**
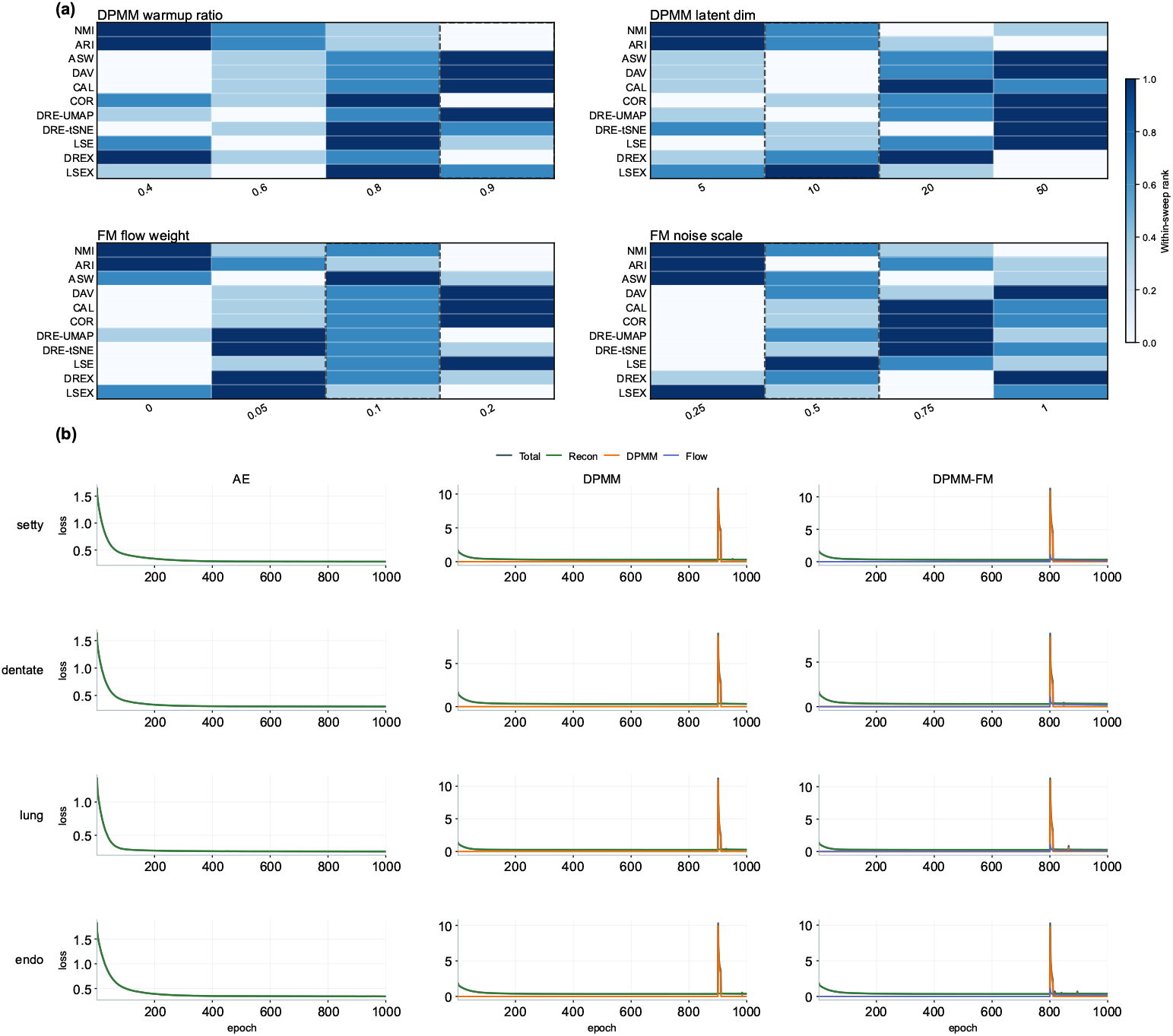
Sensitivity analysis and training dynamics. Panel (a): four 11-metric sensitivity heatmaps summarizing mean-rank performance across the 12-dataset sensitivity suite for DPMM warmup ratio, DPMM latent dimension, FM flow weight, and FM noise scale. Dashed outlines mark default settings. Panel (b): training-loss trajectories on four representative datasets for Pure-AE, DPMM-Base, and DPMM-FM, decomposing total, reconstruction, DPMM, and flow losses over training.

### 3.5. Biological Validation

To assess whether the improved geometry carries biological meaning, we perform perturbation-based gene-importance analysis, latent–gene correlation mapping, UMAP overlay inspection, and Gene Ontology (GO) enrichment [26] on three representative datasets (setty, endoderm, dentate gyrus).

### 3.6 External Benchmark

Figure 5 summarizes the unified 12-dataset external comparison, contrasting the best DPMM variant with 11 representative external baselines—including scVI [2], Cell-BLAST [27], SCALEX [28], *β*-VAE [29], and the MoCo-style CLEAR baseline—across the full 41-metric panel spanning concordance, geometry, projection fidelity, and latent-structure quality.

**Figure 4.**
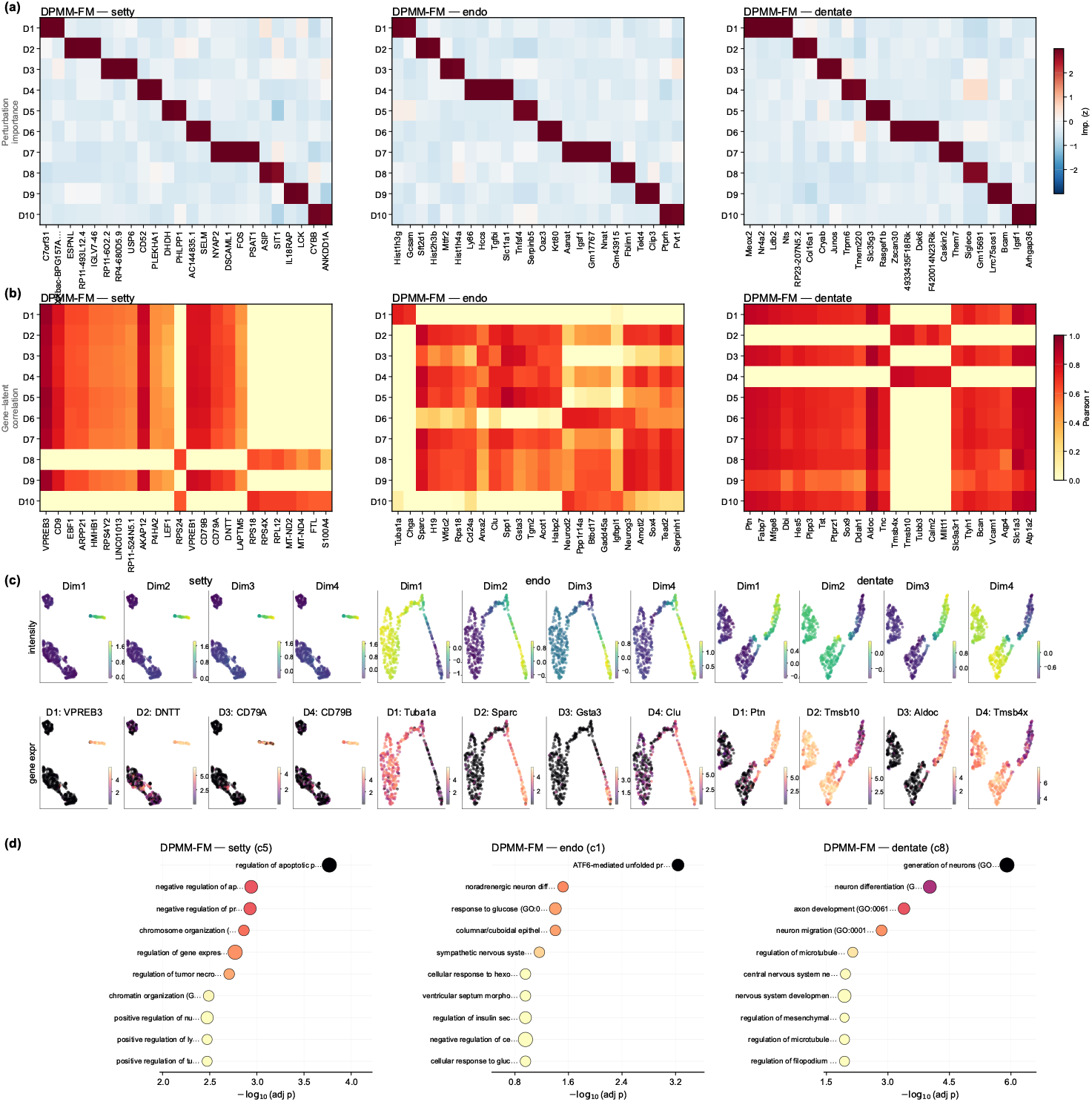
Biological validation. Panel (a): experimental workflow—perturbation-based gene importance analysis on three pre-trained DPMM models (setty, endoderm, dentate gyrus) with forward-pass latent perturbation, top-gene grouping by dominant component, and block-diagonal marker validation. Panel (b): z-scored perturbation importance heatmaps (gene *×* latent component) for DPMM-Base, DPMM-Transformer, and DPMM-Contrastive across all three datasets, with genes sorted by dominant component to reveal block-diagonal structure indicating that each latent dimension captures a distinct expression program.

**Figure 5.**
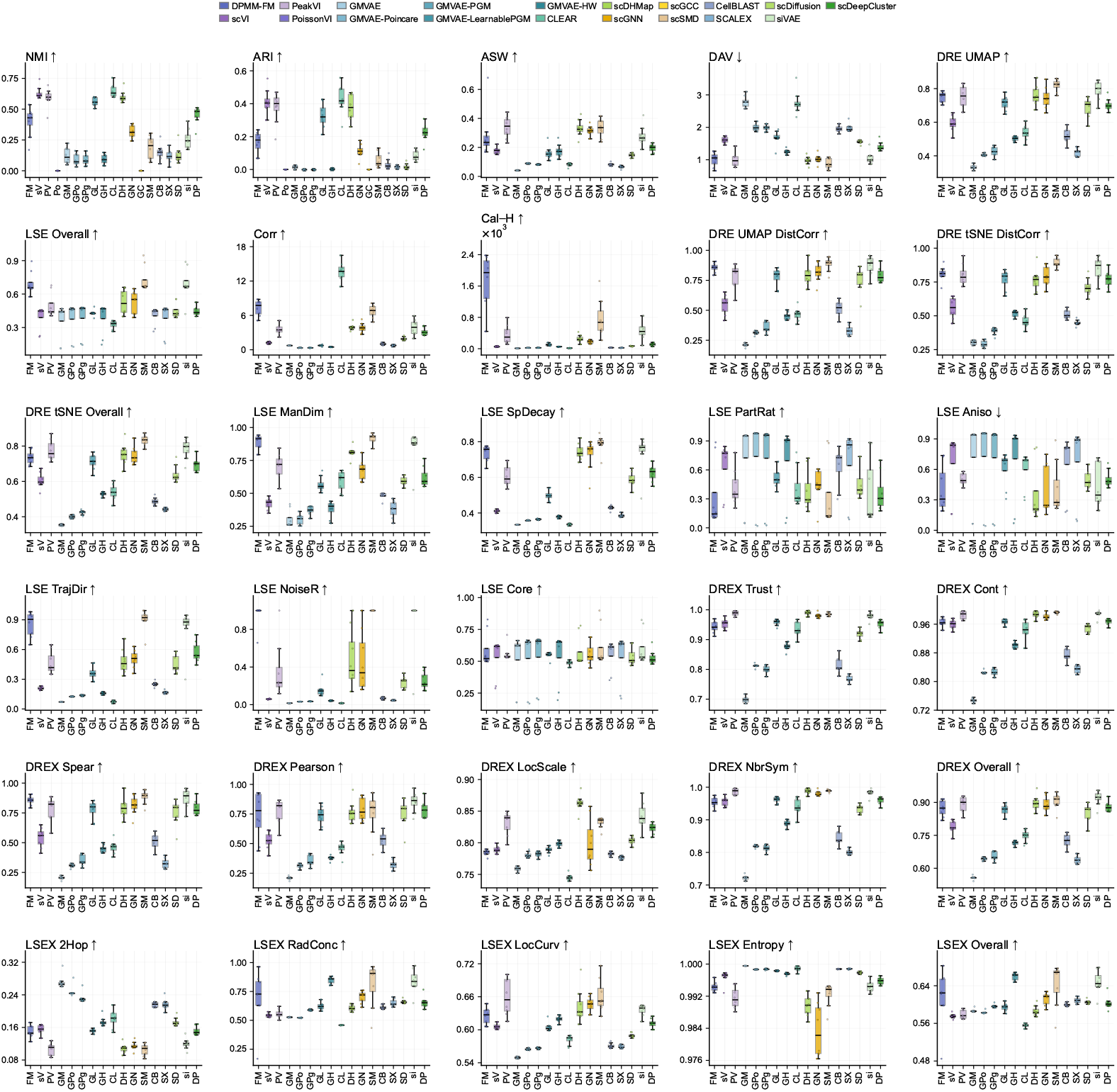
External benchmark. Panel (a): experimental workflow showing 11 representative external baselines evaluated alongside the best DPMM variant under a unified preprocessing pipeline (3k HVGs, log_1_p normalization) across 12 datasets and 41 metrics. Panels (b)–(d): metric boxplot grids for proposed models, classical baselines (PCA, ICA, NMF, FA, Truncated-SVD, UMAP, t-SNE), and deep/probabilistic baselines (scVI [2], scDHA [30], SCALEX [28], *β*-VAE [29], CLEAR, and others), organized by clustering concordance, DRE-UMAP, DRE-tSNE, LSE intrinsic, DREX, and LSEX families.

Table 4 quantifies the external win rates.

**Table 4.** External win-rate summary. Each variant compared against 18 external baselines via Wilcoxon signed-rank tests across 56 datasets. Core win rate = fraction of core-metric comparisons significantly won.

| Model | Total tests | Sig. (all) | Core sig. | Core total | Core win rate |
| --- | --- | --- | --- | --- | --- |
| DPMM-Base | 407 | 176 | 31 | 44 | 0.705 |
| Pure-AE | 407 | 138 | 28 | 44 | 0.636 |

DPMM-Base achieves a core win rate of 70.5%, winning 31 of 44 core-metric comparisons against external baselines, with 176 total significant wins across all metrics. This advantage is concentrated on ASW and DAV; on NMI alone, several external baselines— notably scVI [2] and CLEAR—match or exceed DPMM-Base on individual datasets, consistent with the geometry–concordance trade-off observed internally.

### 3.7. Downstream Classification

Table 5 reports *k*NN classification accuracy and macro-F1 as a supervised probe of representation quality.

**Table 5.**
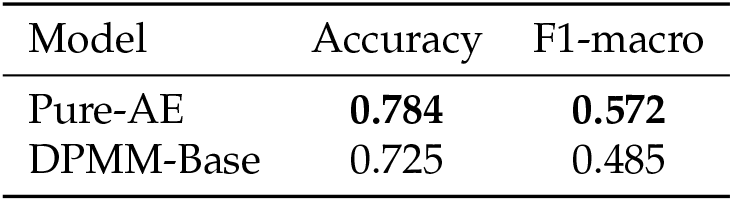
*k*NN downstream classification. Mean accuracy and macro-F1 across 12 datasets. Best value per column in bold.

Pure-AE outperforms DPMM-Base on both accuracy (0.784 vs. 0.725, −7.5%) and macro-F1 (0.572 vs. 0.485, −15.2%). This result is expected and important: *k*NN classification rewards local neighborhood purity, which aligns with label concordance rather than global cluster compactness. It confirms that the DPMM prior’s geometry gains do not translate to better supervised discrimination and that downstream classification tasks should use prior-free representations unless manifold-level structure is specifically required.

### 3.8. Variant Summary

Table 6 ranks the two primary models by composite score.

**Table 6.** Variant ranking. Models ranked by composite score across 56 datasets. Best value per column in bold.

| Model | NMI | ARI | ASW | DAV ↓ | Score |
| --- | --- | --- | --- | --- | --- |
| DPMM-Base | 0.506 | 0.320 | **0.374** | **0.868** | **0.400** |
| Pure-AE | **0.609** | **0.406** | 0.165 | 1.624 | 0.393 |

The composite advantage of DPMM-Base over Pure-AE is small (0.400 vs. 0.393) because the score weights concordance at two-thirds. If geometry alone were the criterion, the gap would be far larger. This observation reinforces the paper’s central claim: the DPMM prior is a geometry-specialized tool, not a general-purpose improvement.

### 3.9. Computational Efficiency

Table 7 reports the computational cost.

**Table 7.** Runtime analysis. Mean training time, throughput, peak GPU memory, and parameter count across 56 datasets. Best value per column in bold.

| Model | Time (s) | Sec/Epoch | GPU (MB) | Params |
| --- | --- | --- | --- | --- |
| Pure-AE | **46.4** | **0.0464** | **922.4** | **1,615,430** |
| DPMM-Base | 50.5 | 0.0505 | 1025.5 | 1,615,430 |

DPMM-Base adds minimal overhead over Pure-AE (50.5s vs. 46.4s, +8.8%), confirming that the DPMM prior’s online refitting is efficient.

## 4. Discussion

### The geometry–concordance trade-off is real and consequential

The central finding of this study is not that DPMM-Base “outperforms” Pure-AE in a universal sense—the composite margin is only 1.8%—but that the two models occupy qualitatively different operating regimes. Pure-AE produces latent spaces that are easy to cluster into ground-truth labels (NMI 0.609, ARI 0.406, *k*NN accuracy 0.784) but geometrically diffuse (ASW 0.165, DAV 1.624). The DPMM prior reverses this profile: compact, well-separated clusters (ASW 0.374, DAV 0.868) whose boundaries do not always align with annotated cell-type labels. This distinction matters practically. For cell-type classification, Pure-AE is preferable. For trajectory analysis, pseudotime inference, or manifold visualization—where global geometric coherence matters more than *K*-means label recovery—DPMM-Base provides a stronger foundation.

### Flow refinement extends the axis, not the optimum

DPMM-FM does not improve upon DPMM-Base across the board; instead, it opens a third operating point. By smoothing the latent manifold via conditional optimal-transport flow [11,12], DPMM-FM achieves the highest projection-fidelity scores (DRE 0.751, LSE 0.695, DREX 0.873) among all variants, but at the cost of lower ASW (0.288 vs. 0.449) and sharply reduced concordance (NMI 0.397, ARI 0.165). The flow field trades local cluster tightness for global manifold smoothness. This makes DPMM-FM the variant of choice when the downstream task is UMAP/t-SNE visualization or distance-preserving embedding, and not when the task is cluster assignment. The three models—Pure-AE, DPMM-Base, DPMM-FM—thus trace a Pareto front from concordance through geometry to projection fidelity, and the correct choice depends on the biological question.

### Why concordance drops: a mechanistic note

The NMI and ARI losses under the DPMM prior are not surprising when examined mechanistically. The online Bayesian Gaussian Mixture refits every 10 epochs [9], pulling latent points toward *its* inferred cluster centers, which may merge or split cell-type boundaries differently from the ground-truth annotation. The result is compact clusters that are internally homogeneous but that may group sub-populations of different annotated types when the DPMM infers that the latent evidence supports fewer (or different) partitions. This is a feature of nonparametric inference, not a bug, but it does reduce concordance with fixed label sets.

### External positioning

The 70.5% core win rate against 18 baselines [2,14,27–30] is driven primarily by geometry metrics. On NMI alone, scVI and CLEAR match or exceed DPMM-Base on several datasets. This is consistent with the trade-off: methods optimized for reconstruction and label recovery (scVI, *β*-VAE) naturally score higher on concordance, while DPMM-Base dominates on compactness and separation. Notably, the concurrent scDAC approach [10], which also couples an autoencoder with a DPMM prior, prioritizes concordance optimization; our work complements it by explicitly quantifying the geometry– concordance cost surface.

### Biological grounding

Perturbation analysis, latent–gene correlations, and GO enrichment [26] on three representative datasets confirm that geometry-improved components capture coherent biological programs (Fig. 4). Importantly, this validates the DPMM prior’s partitioning as biologically meaningful even when it diverges from annotated labels—the clusters correspond to expression programs, not noise.

### Limitations and scope

The concordance cost is the primary limitation: when celltype labeling accuracy is the sole objective, the DPMM prior is counterproductive, and Pure-AE or a dedicated classifier should be preferred. The 90% warmup ratio delays prior activation to near the end of training, raising the question of whether the DPMM is operating on a sufficiently plastic latent space. The two additional hyperparameters in DPMM-FM (flow weight, noise scale) add complexity, though sensitivity analysis (Fig. 3) shows broad robustness. Biological validation covers only three datasets; broader tissue-type coverage would strengthen the generalizability claim. Finally, this study evaluates only feedforward autoencoders; extending the DPMM prior to variational or graph-based encoders [6] remains future work.

## 5. Conclusions

This study systematically characterizes the effect of imposing a Dirichlet Process Mixture Model prior [7,8] on autoencoder latent spaces for single-cell transcriptomics. The core finding is a reproducible geometry–concordance trade-off: across 56 datasets, the DPMM prior improves cluster compactness (ASW +127%) and separation (DAV −47%) while reducing label-recovery metrics (NMI −17%, ARI −21%) and downstream *k*NN accuracy (−7.5%). A second-stage flow refinement (DPMM-FM) [11] extends the axis toward projection fidelity without recovering concordance. The three resulting models— Pure-AE, DPMM-Base, DPMM-FM—form a Pareto front from label recovery through cluster geometry to manifold visualization, and biological validation confirms that geometry-improved components capture coherent gene-expression programs [26]. Rather than proposing the DPMM prior as a universal replacement for prior-free autoencoders, we recommend it as a task-appropriate tool: use DPMM-Base when the downstream analysis requires tight, biologically coherent clusters (trajectory inference [13], spatial organization), and use prior-free models when cell-type classification accuracy is paramount.

## Author Contributions

Z.F. conceived the study, developed the methodology, implemented the software, performed all experiments and analyses, and wrote the manuscript.

## Data Availability Statement

All scRNA-seq datasets used in this study are publicly available. Processed data and code are available at https://github.com/PeterPonyu/PanODE-DPMM.

